# A critical role for PLCG1 in RAS activation by BCR-ABL1 and FLT3-ITD

**DOI:** 10.1101/2023.04.07.536070

**Authors:** May Yee Szeto, Anjeli Mase, Kayla Kulhanek, Saikat Banerjee, Ignacio Rubio, Jeroen Roose, Neil Shah

## Abstract

Myeloid leukemias are frequently associated with pathologically activating mutations in tyrosine kinases [BCR-ABL1 in chronic myeloid leukemia (CML); FLT3 juxtamembrane internal tandem duplication (ITD) mutations, FLT3 and KIT activation loop mutations in acute myeloid leukemia (AML)]. Mutations in these kinases activate RAS, which initiates multiple downstream signaling pathways that regulate cell proliferation, differentiation, and apoptosis. The mechanisms whereby RAS is activated by these kinases is incompletely understood, and a better understanding of the molecular mediators involved in RAS activation may uncover new therapeutic strategies. Here we identify a biologically and therapeutically important novel mechanism whereby BCR-ABL1 and FLT3-ITD activate the critical downstream effector RAS in part through phospholipase C gamma-1 (PLCG1). PLCG1 knockout decreases proliferation of CML and FLT3-ITD-expressing AML cells, reduces RAS nucleotide exchange factor activity, and increases sensitivity of CML cells to BCR-ABL1 tyrosine kinase inhibitors (TKIs). Collectively, these studies suggest that PLCG1 inhibition may augment clinical responses to BCR-ABL1 and FLT3 TKIs in CML and AML.

## Introduction

Clinical experience with BCR-ABL1 and FLT3 tyrosine kinase inhibitors (TKIs) for the treatment of chronic myeloid leukemia (CML) and acute myeloid leukemia (AML) provides compelling evidence for the phenomenon of oncogene addiction, whereby activated signaling molecules become critically important for cell survival^1,2^. Previous studies in our lab have demonstrated that BCR-ABL1-mediated oncogene addiction is facilitated by persistent high levels of MEK-dependent negative feedback, which mitigates BCR-ABL1-dependent RAS activation^3^. Additionally, activating RAS mutations have been identified in CML patients who develop imatinib resistance in the absence of resistance-conferring secondary BCR-ABL1 kinase domain mutations^4^, and in AML patients who develop resistance to gilteritinib in the absence of secondary kinase domain mutations in FLT3-ITD^5^. In the setting of blast phase CML patients, our lab has identified activating RAS mutations that contribute to overt off-target resistance to BCR-ABL1 inhibitors (Reyes et al, in preparation). Together, these observations highlight the key importance of regulating RAS activity in the setting of CML and AML. The mechanism(s) by which BCR-ABL1 and FLT3-ITD activate RAS, a critical effector of downstream signaling, are incompletely understood. MEK inhibition of BCR-ABL1-expressing cells yields robust RAS-GTP levels, thereby providing a tractable, disease-relevant model to address this question in this and other genetic contexts.

RAS is a guanine nucleotide-binding protein that cycles between a GTP-loaded active state and a GDP-loaded inactivate state. RAS-GTP activates various cellular pathways, including the RAF/MEK/ERK and the PI3K/AKT signaling pathways which are involved in many cellular responses such as proliferation, differentiation, and apoptosis^6^. Guanine nucleotide exchange factors (GEFs) help with GTP binding onto RAS whereas GTPase activating proteins (GAPs) promote the hydrolysis of RAS-GTP into RAS-GDP^6^. BCR-ABL1 has previously been shown to activate the RAS/MAPK signaling pathway in part via recruitment of the adaptor protein growth factor receptor-bound protein 2 (GRB2) to a tyrosine-phosphorylated site, Y177, in the N-terminal portion of BCR^7^. Phosphorylation of Y177 in BCR is necessary for BCR-ABL1 - mediated leukemogenesis and mutation of Y177 to phenylalanine (Y177F) abolishes GRB2 binding and diminishes BCR-ABL1 - induced RAS activation assessed by a RAS-responsive element promoter transcription assay in NIH3T3 cells^6^. Subsequent work demonstrated that GRB2 associates with son of sevenless homolog 1 (SOS1), a RasGEF ubiquitously expressed in mammalian cells, to mediate RAS activation^6,8^. Interestingly, another study conducted in BaF3 and 32D myeloid cell lines stably expressing Y177F BCR-ABL1 demonstrated that although GRB2 binding was abolished, RAS activation was not affected^9^. Together, the available data suggest that BCR-ABL1 may recruit multiple signaling molecules to induce RAS activation and the exact mechanisms by which BCR-ABL1 activates RAS are incompletely understood. FLT3-ITD is a receptor tyrosine kinase that localizes both to the plasma membrane (PM) and intracellular endoplasmic reticulum/Golgi compartment of cells and both the PM and intracellular pools of FLT3-ITD can activate downstream signal transduction^10^. FLT3-ITD has been shown to activate RAS at the plasma membrane, whereas the intracellular pool of FLT3-ITD appears incapable of activating RAS in endomembranes^11^. Additionally, studies have shown that the protein tyrosine phosphatase SHP2 binds to FLT3 at phospho-tyrosine residue 599, which is located in the juxtamembrane domain of FLT3 and is duplicated in FLT3-ITD, and is possibly an adapter protein for GRB2 which can associate with SOS1 to mediate RAS activation^12–14^. The exact mechanism of how FLT3-ITD activates RAS remains unknown.

Phospholipase C (PLC) enzymes catalyze the formation of the second messengers inositol 1,4,5-triphosphate (IP3) and diacylglycerol (DAG) from phosphatidylinositol 4,5-bisphosphate (PIP2), which activate intracellular calcium and protein kinase C (PKC) signaling pathways as well as GEFs of the RasGRP family^15,16^. Since the p190 isoform of BCR-ABL1 (p190-BCR-ABL1), which is more commonly found in BCR-ABL1+ acute lymphoblastic leukemia, has been reported to interact with PLCG1, and ABL1 has been demonstrated to interact directly with PLCG1^17,18^, we sought to determine if and how the p210 isoform of BCR-ABL1, which is present in ∼99 percent of CML cases, utilizes the PLCG1 signaling pathway to activate RAS, and to extend our findings to FLT3-ITD. Additionally, we sought to identify the potential relevance of a PLCG1 signaling axis to CML and AML cellular proliferation and TKI sensitivity. Since RAS is a critical effector of BCR-ABL1- and FLT3-ITD-mediated downstream signaling, and activating mutations in RAS have been identified in CML and AML patients who are resistant to BCR-ABL1 or FLT3 TKI therapy, a better understanding of how BCR-ABL1 and FLT3-ITD activate RAS may help develop novel therapeutic strategies.

## Results

### BCR-ABL1 and FLT3-ITD directly activate the PLCG1 signaling axis and BCR-ABL1 physically associates with PLCG1 in a kinase activity independent manner

We first evaluated whether there is an interplay between the kinase activity of BCR-ABL1 or FLT3-ITD and the PLCG1 signaling axis with the use of relatively selective kinase inhibitors. Inhibition of BCR-ABL1 kinase activity with imatinib or dasatinib in the CML cell line K562 abolished phosphorylation of PLCG1 at tyrosine 783, which is critical for PLCG1 activation and is accepted as a marker for PLCG1 activity^19,20^ (**Figure 1a**). Similarly, inhibition of FLT3-ITD kinase activity with AC220 (quizartinib) in the FLT3-ITD-dependent AML patient-derived cell line Molm14 eliminated phosphorylation of PLCG1 (**Figure 1b**). Since p190-BCR-ABL1 has been reported to interact with PLCG1 and we found that inhibition of BCR-ABL1 activity reduced activation of PLCG1, this raised the question of whether the p210 isoform of BCR-ABL1 can directly activate PLCG1. By co-immunopreciptation, we identified a BCR-ABL1 - PLCG1 complex in K562 cells in both the presence and absence of BCR-ABL1 kinase activity (**Figure 1c**). Notably, a cABL1-PLCG1 complex was also detected in a kinase independent manner (**Figure 1c**). Together, these data suggest that BCR-ABL1 and ABL1 can directly bind to PLCG1, and that BCR-ABL1 can phosphorylate PLCG1 on tyrosine 783. Moreover, ABL1 sequences appear to be sufficient for PLCG1 binding.

**Figure 1.**
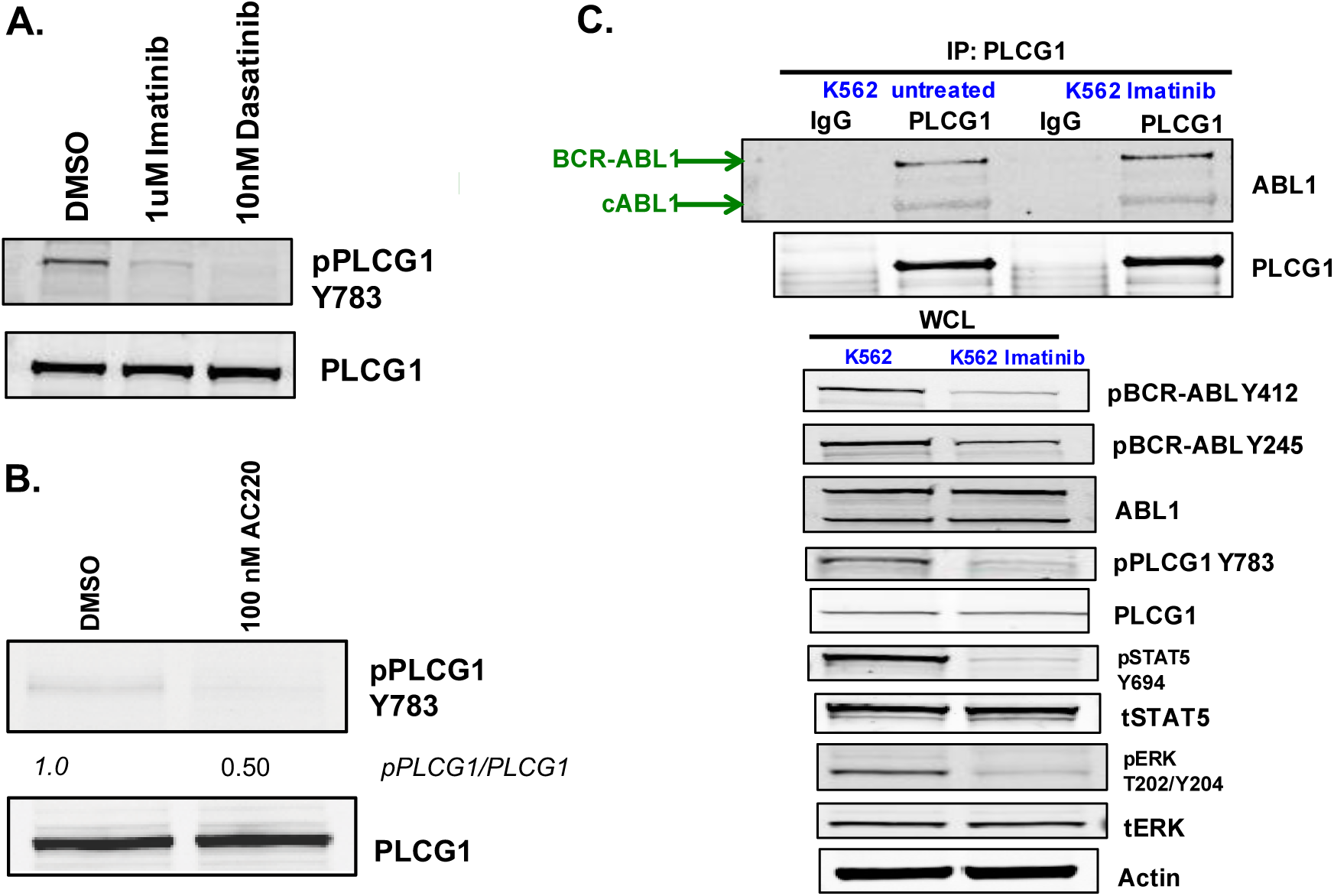
BCR-ABL1 and FLT3-ITD directly activate the PLCG1 signaling axis and BCR-ABL1 physically associates with PLCG1 in a kinase activity independent manner. **A.** BCR-ABL1 inhibition decreases PLCG1 activity. Western immunoblot analysis of PLCG1 activity in K562 cells treated with indicated drugs for 1h. **B.** FLT3-ITD inhibition decreases PLCG1 activity. Western immunoblot analysis of PLCG1 activity in Molm14 cells treated with indicated drugs for 1h. **C.** PLCG1 physically associates with BCR-ABL1 and ABL1 in a kinase independent manner. Endogenous PLCG1 was pulled down in K562 cells following 1h treatment with 0.1% DMSO or 1uM Imatinib. Coimmunoprecipitated ABL was blotted for (top) and whole cell lysates were blotted for (bottom).

### PLCG1 activates RAS in BCR-ABL and FLT3-ITD expressing cells

We next assessed whether the PLCG1 signaling axis is involved in BCR-ABL1 - and FLT3-ITD - mediated RAS activation. Previous work in our lab has demonstrated that BCR-ABL1-mediated oncogene addiction is facilitated by persistent high levels of MEK-dependent negative feedback, which mitigates BCR-ABL1 dependent RAS activation^3^. As previously published work on RAS activation by BCR-ABL1 has yielded inconsistent findings in different cellular contexts, we utilized CML patient-derived cells. The K562 cell line, derived from a myeloid blast crisis CML patient, has low basal RAS-GTP levels. As demonstrated previously (Asmussen et al), treatment with the allosteric MEK inhibitor PD0325901 for 12 hours yields robust RAS-GTP levels, whereas co-treatment with imatinib/PD0325901 abolishes RAS-GTP loading, demonstrating that RAS activation following MEK inhibition in CML cells is mediated by BCR-ABL1 kinase acitivity. We sought to utilize the robust RAS-GTP levels observed following MEK inhibition in K562 cells as a tractable, disease-relevant model to address the question of how BCR-ABL1 activates RAS. We assessed whether the PLCG1 signaling axis is involved in BCR-ABL1 mediated RAS activation through chemical and genetic approaches. Following treatment with a MEK inhibitor (PD0325901), the PLC inhibitor U73122 reduced RAS-GTP levels by about 50% compared to vehicle. These findings were confirmed in KU812 cells, an independent myeloid blast crisis CML patient-derived cell line (**Figure 2a**). Since the PLC inhibitor U73122 is nonspecific, we next took a genetic approach to knockout PLCG1 in K562 cells using clustered regularly interspaced short palindromic repeats (CRISPR) technology. Two different guide RNA sequences were used to knockout PLCG1 by CRISPR^21^, which abolished PLCG1 protein levels but did not affect PLCG2 protein levels (**Figure 2b and Supplementary Figure 1**). Knockout of PLCG1 resulted in decreased basal ERK activity levels compared to parental control cells (**Figure 2b**). Genetic deletion of PLCG1 in two independent clones expressing different guide RNA sequences for PLCG1 was associated with RAS-GTP levels of approximately 50% following MEK inhibition as compared to parental cells (**Figure 2b**). Taken together, these data suggest that PLCG1 contributes approximately half of the RAS activation in BCR-ABL1-expressing human myeloid cells. Similarly, treatment of FLT3-ITD expressing Molm14 cells with the MEK inhibitor PD0325901 yields robust RAS-GTP levels, demonstrating the presence of high levels of MEK-dependent negative feedback in this context as well. Co-treatment with AC220/PD0325901 prevented RAS-GTP loading (**Figure 2c)**, demonstrating that RAS activation in these cells is facilitated by FLT3-ITD kinase activity. Upon co-treatment with a MEK and PLC inhibitor, FLT3-ITD expressing cells have reduced RAS-GTP levels compared to vehicle (**Figure 2c**). This data suggests that FLT3-ITD-mediated RAS activation is achieved in part through activation of PLCG1 in FLT3-ITD driven AML.

**Figure 2.**
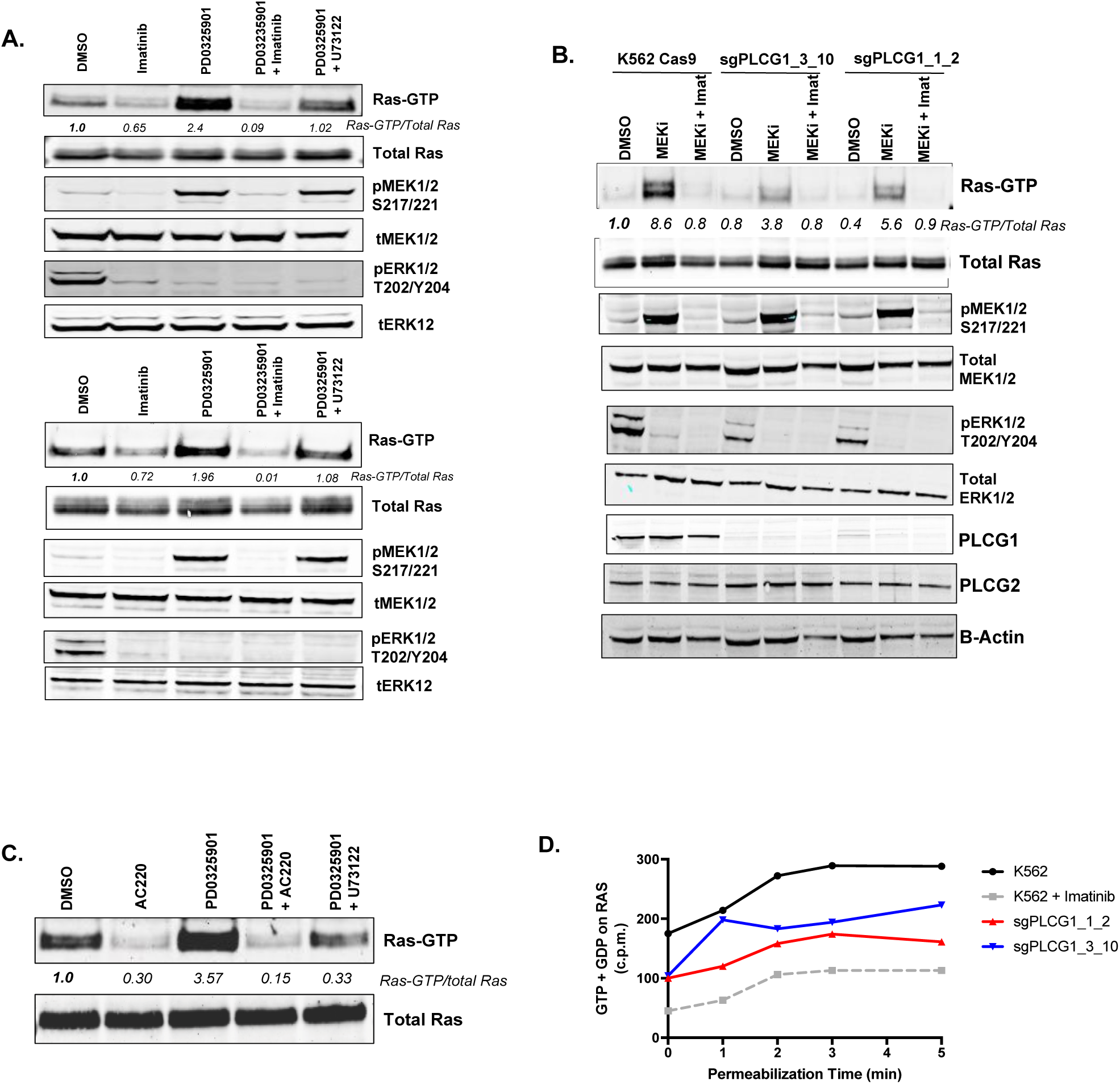
PLCG1 activates RAS in BCR-ABL and FLT3-ITD expressing cells. **A.** PLC inhibition diminishes RAS activation in BCR-ABL1 expressing cells. Western immunoblot analysis of RAS, MEK, and ERK activity in K562 cells (top panel) and KU812 cells (bottom panel) that were treated with 0.1% DMSO, 1uM Imatinib, 500nM PD0325901, or Imatinib and PD0325901, or Imatinib and 5uM U73122 for 12 hours. **B.** PLCG1 is responsible for RAS activation in BCR-ABL1 expressing cells. Western immunoblot analysis of RAS, MEK, and ERK activity in K562 cells with PLCG1 knocked out treated with DMSO, MEK inhibitor (500nM PD0325901), or MEK inhibitor and Imatinib (1uM) for 12 hours. **C.** PLC inhibition diminishes RAS activation in FLT3-ITD expressing cells. Western immunoblot analysis of RAS activity in Molm14 cells that were treated with 0.1% DMSO, 100nM AC220, 500nM PD0325901, or AC220 and PD0325901, or AC220 and 5uM U73122 for 12 hours. **D.** PLCG1 knockout BCR-ABL1 expressing cells have reduced RAS nucleotide exchange activity. Accumulation of radiolabeled guanine nucleotide in counts per minute that was loaded onto RAS in the basal state over time as measured with the digitonin exchange assay. K562 treated with 1uM Imatinib for 12h was used as a control. Shown is representative data from one of three different independent experiments performed.

To investigate how loss of PLCG1 in BCR-ABL1 expressing cells directly affects the equilibrium between RAS-GDP/RAS-GTP, we took advantage of a RAS nucleotide exchange assay to evaluate the rate of RAS-GTP loading^22^. Whereas traditional RAS pulldown only provides an assessment of net RAS-GTP levels, this assay is sensitive and can dissect the rate of nucleotide exchange guanine nucleotide exchange factors^23^. Radioactivity levels increased quickly and were saturated by 3 minutes in K562 cells, indicating that all RAS molecules had undergone one round of nucleotide exchange (**Figure 2d**). Treatment with imatinib caused a marked reduction of nucleotide uptake by RAS, suggesting that the RAS-GTP loading in K562 cells is dependent on BCR-ABL1 kinase activity (**Figure 2d**). In both K562 PLCG1 knockout clones, a marked reduction (approximately 50%) of nucleotide uptake by RAS was observed (**Figure 2d**). This suggests that PLCG1 knockout cells have reduced nucleotide exchange factor activity. Given the observation that control K562 cells have similar steady-state levels of RAS-GTP compared to PLCG1 knockout cells (**Figure 2a**), the data argues for a high basal RasGAP activity in control K562 cells, and suggest that the decreased RAS-GTP levels we observed with the traditional RAS pulldown assays following MEK inhibition in the two PLCG1 knockout clones is due to reduced nucleotide exchange factor activity. Collectively, these data suggest that in BCR-ABL1 expressing cells there is constitutive loading and hydrolysis of GTP on RAS, with the former process being dependent in part on PLCG1.

### Knockout of PLCG1 decreases proliferation of BCR-ABL1 and FLT3-ITD expressing cells and modulates responsiveness to BCR-ABL1 TKIs but not to FLT3 TKIs

To address the biological importance of PLCG1 toward BCR-ABL1 and FLT3-ITD expressing cells, cell growth competition assays were performed. Parental K562 and Molm14 cells (mCherry negative) were mixed in a 1:1 ratio with either PLCG1 knockout cells or cells expressing non-targeting control guide RNA (mCherry positive; conferred by guide RNA expression), and mCherry expression was assessed over time. Whereas cells expressing control guide RNA grew with similar kinetics to parental K562 or Molm14 cells (and the proportion of mCherry-positive cells remained approximately 50% over time), PLCG1 knockout BCR-ABL1 or FLT3-ITD cells displayed a slower growth rate, as evidenced by a decrease in mCherry-positive cells over time (**Figures 3a and 3b**). K562 and Molm14 cells expressing control guide RNA alone or PLCG1 knockout cells alone remained 100% mCherry positive during the experiment, and parental K562 and Molm14 cells alone remained 100% mCherry negative (data not shown). These data suggest that PLCG1 contributes to the proliferation of CML and AML cells.

**Figure 3.**
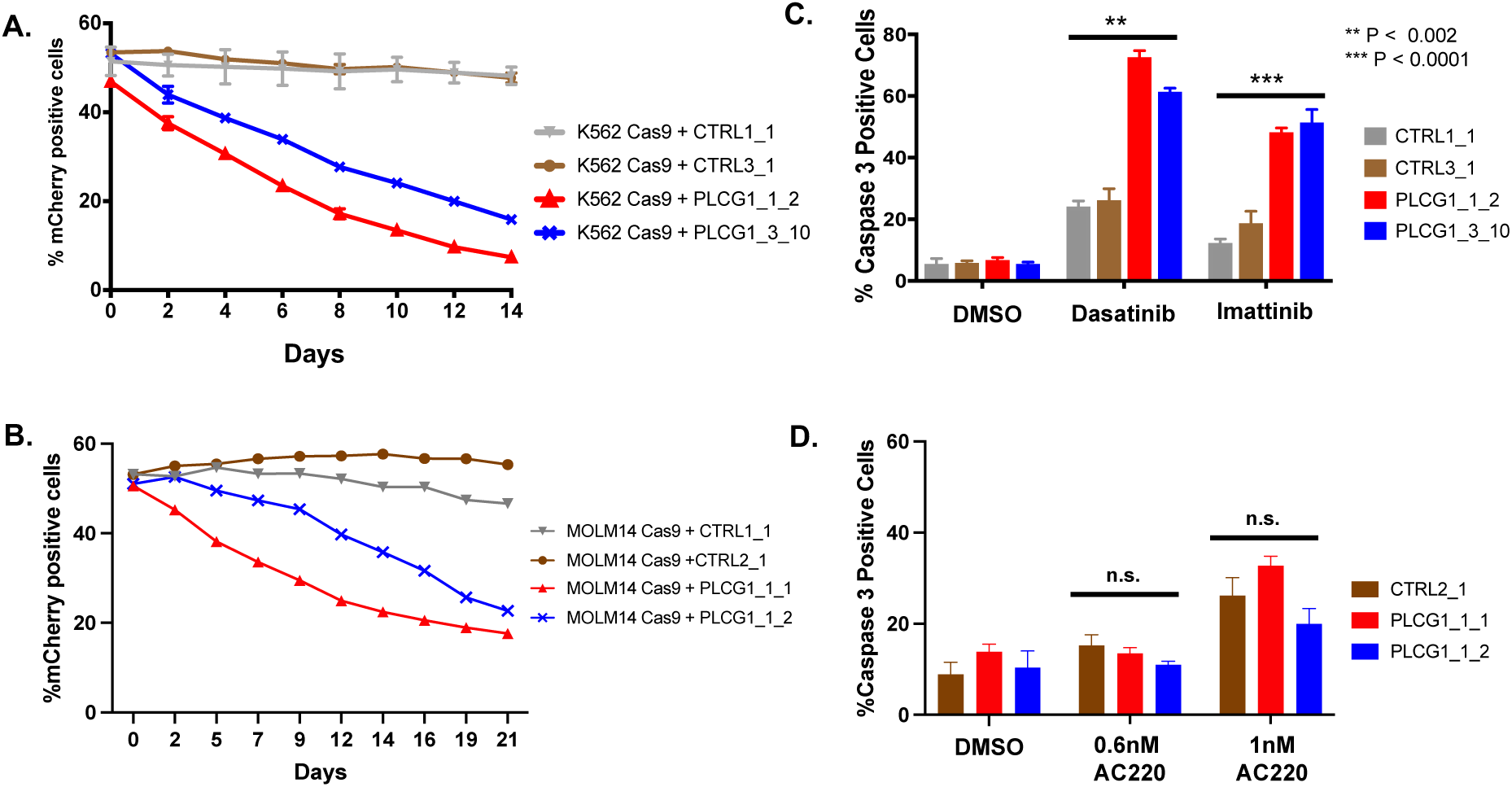
Knockout of PLCG1 decreases proliferation of BCR-ABL1 and FLT3-ITD expressing cells and modulates responsiveness to BCR-ABL1 inhibitors but not to FLT3 inhibitors. **A.** Knockout of PLCG1 decreases growth rate of BCR-ABL1 expressing cells. Equal number of K562 Cas9 cells (mCherry negative) and control or PLCG1 knockout cells (mCherry positive) were plated and mCherry expression was assessed over time. **B.** Knockout of PLCG1 decreases growth rate of FLT3-ITD expressing cells. Equal number of Molm14 Cas9 cells (mCherry negative) and control or PLCG1 knockout cells (mCherry positive) were plated and mCherry expression was assessed over time. **C.** Knockout of PLCG1 in BCR-ABL1 expressing cells increases sensitivity to BCR-ABL1 tyrosine kinase inhibitors. Flow cytometric detection of cleaved caspase-3 in control and PLCG1 knockout BCR-ABL1 expressing cells treated with 0.1% DMSO, 0.4nM Dasatinib, or 200nM Imatinib for 48h. P-values were calculated with two tailed unpaired *t*-test. **D.** Knockout of PLCG1 in FLT3-ITD expressing cells does not modulate sensitivity to FLT3 tyrosine kinase inhibitors. Flow cytometric detection of cleaved caspase-3 in control and PLCG1 knockout FLT3-ITD expressing cells treated with 0.1% DMSO, 0.6nM AC220, or 1nM AC220 for 48h. P-values were calculated with two tailed unpaired *t*-test.

Since deletion of PLCG1 negatively impacts RAS activation and decreases the growth rate of CML and AML cells, we next assessed whether the PLCG1 signaling axis impacts BCR-ABL1 and FLT3 tyrosine kinase inhibitor sensitivity. PLCG1 knockout CML cells demonstrate increased sensitivity to imatinib and dasatinib compared to parental cells by assessing cell proliferation using a Cell Titer Glo assay (**Supplementary Figure 2a**). We also observed that following treatment with imatinib or dasatinib that PLCG1 knockout BCR-ABL1 expressing cells commit to apoptosis to a far greater extent than cells expressing control guide RNA (**Figure 3c, Supplementary Figure 2b**). Together, these data highlight the functional role of the PLCG1 signaling axis in the survival and proliferation of BCR-ABL1-expressing cells. Interestingly, while knockout of PLCG1 decreases proliferation of FLT3-ITD expressing cells, we did not observe increased sensitivity to the FLT3 tyrosine kinase inhibitor AC220 (**Figure 3d**).

The observation that deletion of PLCG1 increases sensitivity to BCR-ABL1 tyrosine kinase inhibitors led us to next determine whether combining PLC inhibitor and BCR-ABL1 tyrosine kinase inhibitors may have an additive or synergistic effect in reducing cell viability. To explore this possibility, K562 cells were treated with various concentrations of the PLC inhibitor U73122 (IC50 determined to be 5uM in K562 cells with 48hrs treatment) with and without various concentrations of imatinib (IC50 determined to be 400nM in K562 cells with 48hrs treatment) and cell viability following drug treatment was assessed with a Cell Titer Glo assay. We observed that the addition of imatinib to U73122 led to decreased cell viability (**Supplementary Figure 2c**). However, when the Bliss model of independence or the Chou Talalay method to assess additivity or synergy were employed to analyze the data, statistical significance was not observed (**Supplementary Figure 2d**). The observations do not rule out the possibility that combining a PLCG1 inhibitor to BCR-ABL1 tyrosine kinase inhibitors may have an additive or synergistic effect, as U73122 is a nonspecific PLC inhibitor. However, sufficiently selective PLC inhibitors to address this question do not currently exist.

## Discussion

While hyperactive RAS signaling is a common feature of numerous malignancies, the mechanism(s) of RAS activation in many settings is incompletely understood. We have identified a biologically and therapeutically targetable pathway whereby BCR-ABL1 and FLT3-ITD activate the critical downstream effector RAS in part through PLCG1 (**Figure 4**). Prior to our work, the only other evidence that PLCG1 regulates the activity of guanine nucleotide exchange factor was in human prostate cancer cells^24^. There have been some studies implicating a possible role of the PLCG1 signaling axis in the BCR-ABL1 context. The first piece of evidence comes from the observation that p190-BCR-ABL1 can physically associate with PLCG1, and PLCG1 is tyrosine phosphorylated in p190-BCR-ABL1 expressing cells^17^. The second comes from studies demonstrating a PLCG1-dependent mechanism of mTOR/p70S6-kinase activation in BCR-ABL1 expressing cells^25^. Our results suggest that BCR-ABL1 may activate PLCG1 by direct phosphorylation, which can lead to activation of RAS through a yet unknown molecular mechanism. We speculate that PLCG1 activates RAS through the RasGRP family of guanine nucleotide exchange factors which contain a diacylglycerol binding domain, as we have found that stimulation of CML cells with phorbol myristate acetate (PMA), a diacylglycerol mimetic, yields robust RAS activation **(Supplementary Figure 3a)**. While there are four members of the RasGRP family of RasGEFs, K562 cells do not express detectable amounts of RasGRP1 or RasGRP3 protein (data not shown) and additional studies have shown that K562 cells only express RasGRP2 and RasGRP4^26^, with the former functioning as a nucleotide exchange factor for the small GTPase Rap^23,27^. We attempted to knockout RasGRP4 in K562 cells with CRISPR but were only able to generate K562 cells that were heterozygous for RasGRP4 (**Supplementary Figure 3b**). Loss of one copy of RasGRP4 decreased proliferation of BCR-ABL1 expressing cells (**Supplementary Figure 3c**), but did not affect Ras-GTP basally or following MEK inhibition (**Supplementary Figure 3d**). The difficulties in generating RasGRP4 knockout cells suggests a possible critical biological importance of RasGRP4 in mediating cell survival in CML cells. Interestingly, in the context of AML, a recent study in AML1-ETO-driven AML suggests that PLCG1 is specifically required for leukemic stem cell renewal^28^. The majority of patient samples and cell lines that demonstrated the importance of PLCG1 harbored FLT3-ITD or KIT mutations, but the mechanism of PLCG1 activation was not investigated^28^. Our study is the first to highlight the role of PLCG1 in RAS activation in FLT3-ITD-driven AML cells.

**Figure 4.**
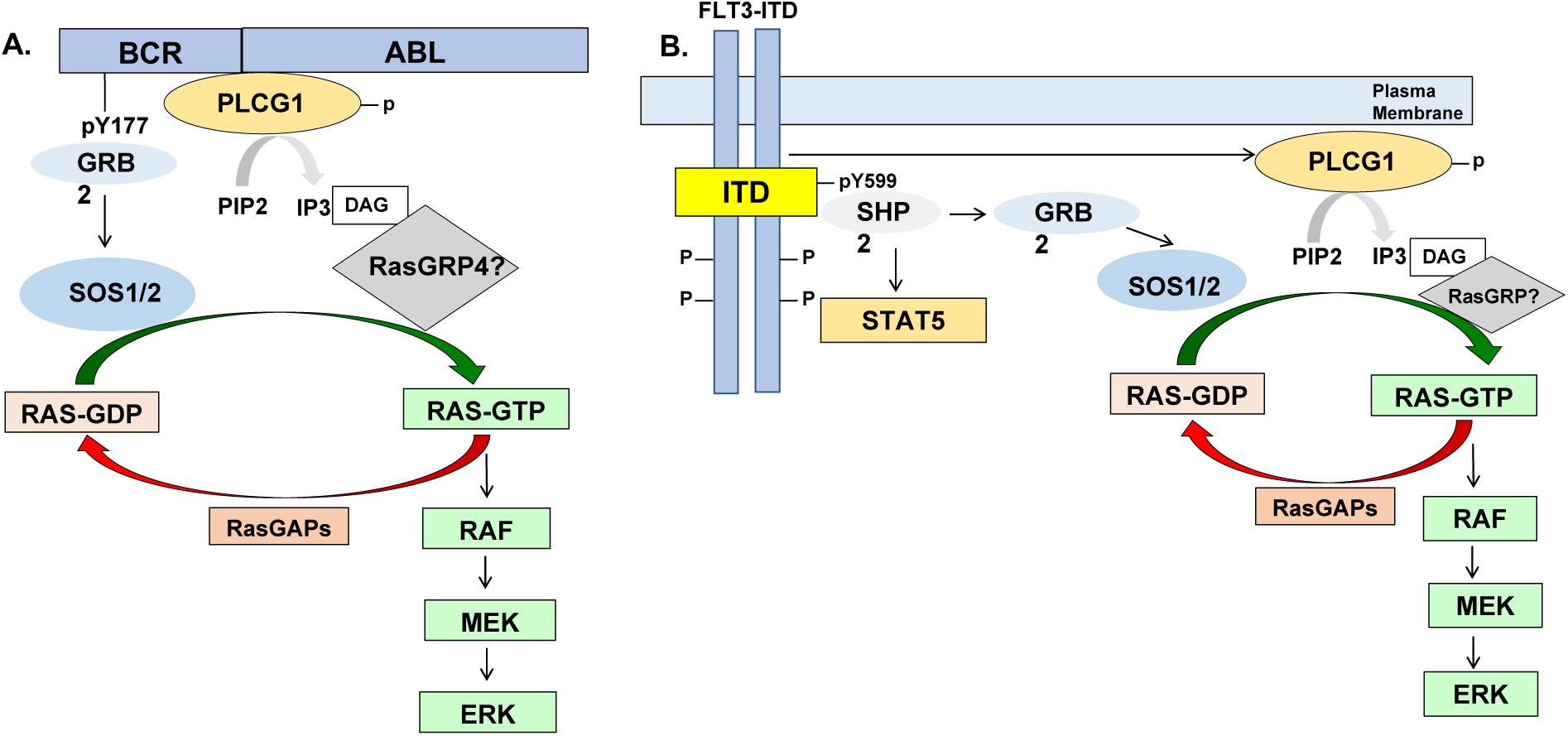
Summary of BCR-ABL1 and FLT3-ITD mediated RAS activation. **A.** Summary of what is known about BCR-ABL1 mediated RAS activation and proposed model of BCR-ABL1 mediated RAS activation. This study highlights a novel mechanism whereby BCR-ABL1 physically associates with PLCG1, which may activate the RasGEF RasGRP4 to activate RAS. **B.** Summary of what is known about FLT3-ITD mediated RAS activation and proposed model of FLT3-ITD mediated RAS activation. This study highlights a novel mechanism whereby PLCG1 partly activates RAS through an unknown mechanism.

The biological relevance of the PLCG1 signaling axis in BCR-ABL1 expressing cells can be appreciated by the observation that loss of PLCG1 either through chemical or genetic means decreases cell proliferation and increases sensitivity to BCR-ABL1 TKIs (**Figure 3**). Notably, some studies have suggested the importance of this signaling pathway in the context of BCR-ABL1. First, inhibiting PLCG1 with a PLC inhibitor or siRNA mediated inhibition of PLCG1 impairs cell proliferation and induces apoptosis in Ba/F3 and 32D cells expressing BCR-ABL1^25^. Additionally, PLCG inhibition in primary mononuclear cells from CML patients or in CML CD34+ hematopoietic progenitor cells induced apoptosis, but not in cells from healthy donors. The degree of apoptosis observed was further increased by the addition of imatinib^25^. Lastly, *in vivo* experiments where mice were injected with Ba/F3 cells expressing p190-BCR-ABL1 and shRNA against PLCG1 had delay in disease onset and increased survival compared to mice expressing p190-BCR-ABL1 and non-targeting shRNA^25^. Notably in other studies, it has been reported that increased phosphorylation of PLCG1 at tyrosine residue 783 (a marker of PLCG1 activity) is associated with primary clinical resistance to BCR-ABL1 tyrosine kinase inhibitors, suggesting that an inability to effectively inhibit this signaling axis may lead to survival of CML stem and early progenitor cells, and thereby contribute to suboptimal therapeutic outcomes^29^. Collectively, our work and these studies suggest an important role of the PLCG1 signaling axis in the survival of BCR-ABL1 expressing cells. Development of potent and selective PLCG1 inhibitors, and investigating their activity in conjunction with BCR-ABL1 TKIs to possibly achieve better cytotoxicity and deeper clinical remissions in CML patients is warranted.

In the context of FLT3-ITD AML, while deletion of PLCG1 negatively impacts RAS activation and decreases the growth rate of cells, there appears to be no impact on FLT3 tyrosine kinase inhibitor sensitivity (**Figures 2 and 3**). Our findings agree with previous studies have shown that RAS activation is essential for the proliferation of FLT3-ITD expressing AML cells^30–32^. The observation that deletion of PLCG1 does not modulate FLT3 TKI sensitivity likely highlights the fact that FLT3-ITD expressing cells may rely more on other signaling cascades like STAT5 in addition to RAS/MAPK for survival^30,33^ and the intrinsic differences between BCR-ABL1 and FLT3-ITD.

In conclusion, we find that BCR-ABL1 and FLT3-ITD activate the critical downstream effector RAS in part through PLCG1. PLCG1 is important for cell proliferation and survival of CML and AML cells. Collectively, our studies suggest that PLCG1 inhibition may augment clinical responses to BCR-ABL1 and FLT3 TKIs in CML and AML.

## Experimental Methods

### Cell Line Propagation and Cell Line Generation

All cell lines were grown in RPMI-1640 medium supplemented with 10% fetal bovine serum (Omega Scientific), 1% L-glutamine and penicillin/streptomycin (Invitrogen).

K562 and Molm14 PLCG1 knockout cells were generated by lentiviral transduction, first with human codon-optimized S. pyogenes Cas9 protein and subsequently superinfected with PLCG1 guide RNA plasmids. Specifically, to generate PLCG1 knockout cell lines, parental K562 or Molm14 cells were first infected with lentivirus for human codon-optimized S. pyogenes Cas9 protein (lentiCas9-Blast plasmid) [33]. Following blasticidin selection, K562 or Molm14 cells stably expressing Cas9 (K562 Cas9, Molm14 Cas9) were infected with lentivirus for guide RNAs against PLCG1. Three different guide RNA sequences targeting PLCG1 were used [24]. PLCG1 guide 1 RNA sequence is ATAGCGATCAAAGTCCCGTG, PLCG1 guide 2 RNA sequence is AGACCCCTTACGAGAGATCG, and PLCG1 guide 3 RNA sequence is CTCAGTGGATCGGAATCGTG. The guide RNA sequences were cloned into MP783 plasmid, a pSico based sgLenti guide RNA library vector after AarI digest, which contains mCherry as well as puromycin marker. Following lentiviral transduction and puromycin selection, K562 Cas9 sgPLCG1 and Molm14 Cas9 sgPLCG1 bulk population cells were subject to limiting dilutions to isolate clones that have PLCG1 knocked out.

To generate K562 RasGRP4 knockout cell lines, a Clustered Regularly Interspaced Short Palindromic Repeats Ribonucleoprotein (CRISPR RNP) approach utilizing two different guide RNAs targeting each gene of interest were used. Guide RNA sequences for each gene of interest were designed such that when two guide RNAs cut, a sizeable deletion (>1000 bp) that results in the generation of a premature stop codon occurs. Guide RNA sequences for RasGRP4 are TCTTCAGCAAGACTTGCCAG and AGAATCACTTGAACCTCAGG. Guide RNAs were ordered from Dharmacon and resuspended in 10mM Tris 150mM NaCl buffer, pH 7.4 to 80uM. The RNP complex was assembled by mixing the two-different guide RNAs for each gene of interest together with tracrRNA (ordered from Dharmacon and resuspended in 10mM Tris 150mM NaCl buffer, pH7.4 to 80 uM). Following incubation for ten minutes at 37°C, human Cas9 protein (ordered from QB3 MAcroLab at 40 uM) was added to the mixture and incubated at 37°C for an additional fifteen minutes. 3uL of the total Cas9-RNP complex was added to 200,000 K562 cells resuspended in 18uL SF cell solution (Lonza) and subjected to nucleofection with the Lonza 4D nucleofectorsystem using manufactured specified conditions for K562 cells. Following nucleofection, cells were allowed to recover for forty-eight hours before a screening PCR was performed on the bulk nucleofected cell population. Sequences for the primers used in the deletion screening reaction are as follows. RasGRP4: 5’- GGGCTGGGTGGAATGAAACT-3’ (forward primer) and 5’-CCCCACCTTCCCTTGAAATGA-3’ (reverse primer) where a successful deletion generates a 877bp product. Following verification that gene editing occurred in the bulk cell population, live single cells were sorted with a FACS Aria II machine into 96 well plates. Screening PCR was performed using the aforementioned primers once the clones grew up. Genomic DNA from cells were isolated by QuickExtract DNA Extraction Solution (Epicenter). Briefly, 5uL cells were added to 10uL QuickExtract DNA Extraction Solution and were incubated at 65 °C for 10 minutes followed by 95 °C for 5 minutes. 2uL of the resulting DNA was used for PCR with Taq polymerase in ThermoPol Buffer per manufacturer protocol (New England BioLabs).

### Inhibitors and Drug Treatments

DMSO stock solutions of imatinib and dasatinib were generated at UCSF. The MEK inhibitor PD0325901, PLC inhibitor U73122, FLT3 inhibitor AC220 were purchased from Selleckchem. PMA was purchased from Sigma-Aldrich. All drug exposures were performed at a cell density of 5 x 10^5^ cells/mL. For RAS-GTP pulldown experiments and PMA stimulation experiments, cells were washed 3x with 1X PBS and plated in media with reduced serum (0.1% fetal bovine serum in RPMI 1640 medium).

### Cell lysis, protein immunoprecipitation, and protein co-immunoprecipitation

For RAS-GTP pulldown experiments, 5×10^6^ cells were lysed for thirty minutes at 4°C in RAS IP buffer (50mM Tris pH 7.5, 125mM NaCl, 6.5mM MgCl_2_, 5% glycerol, 0.2% NP40) supplemented with 1% protease inhibitor cocktail set III and 1% phosphatase inhibitor cocktail set II (Millipore). Following clarification by centrifiguation, cell lysates were tumbled for one hour at 4°C with 20uL of RAS assay reagent (Millipore; #14-278).

For co-immunoprecipitation experiments, 20×10^6^ cells were lysed for thirty minutes at 4°C in 1mL TAP lysis buffer (50mM Tris-HCl pH7.5, 125mM NaCl, 5% glycerol, 0.2% NP40,1.5mM MgCl2) supplemented with 1% protease and 1% phosphatase inhibitors. Following clarification by centrifugation, cell lysates were incubated with 2.5ug anti-PLCG1 antibody (Cell Signaling Technology 2822) or 2.5ug rabbit IgG (Santa Cruz Biotech 2027) or 2.5ug Flag tag antibody (Cell Signaling Technology 8146) or 2.5ug mouse IgG (Santa Cruz Biotech 2025) that was pre-conjugated to 25uL protein G agarose beads (Prometheus #20-537) in a rotating tumbler for forty-five minutes at 4°C. Immunoprecipitates were collected and immunoblotted.

### Antibodies and Western Immunoblot

Cell lysates were resolved by SDS-PAGE and transferred to nitrocellulose membrane as previously described [34]. Flag tag (cat. 8146), PLCG2 (cat. 3872), Beta-actin (cat. 4967), Phospho-specific and total antibodies for MEK1/2 (phospho-Ser 217/221) (cat. 9121 and 9122), ERK1/2 (phospho-Thr202/Tyr204) (cat. 4370 and 9107), PLCG1 (phospho-Tyr783) (cat. 2821 and 2822), cABL (phospho-Y412 and phospho-Y245) (cat. 2865 and 2861), STAT5A/B (phospho-Tyr694) (cat. 9351) were purchased from Cell Signaling Technology. Antibodies against total cABL (Ab-3) (cat. OP20) and total RAS (cat. 05-0516) were purchased from Millipore. Antibody for GAPDH (cat. Sc-25778) were purchased from Santa Cruz Biotechnology. Odyssey imaging technology and software was used for western immunoblot visualization and quantification (Licor).

### Competition growth assay

PLCG1 knockout cells: A total of 0.2×106 cells were plated in a 1mL final volume in a 24 well plate. K562 Cas9 or Molm14 Cas9 cells (mCherry negative), K562 sgCTRL or Molm14 sgCTRL clonal cells (mCherry positive due to guide RNA expression), K562 PLCG1 or Molm14 PLCG1 knockout clonal cells (mCherry positive due to guide RNA expression) were plated alone to monitor mCherry expression of logarithmic growing cells over time. An equal number (0.1×10^6^ cells) of K562 Cas9 and K562 PLCG1 knockout cells or control cells, and Molm14 Cas9 and Molm14 PLCG1 knockout cells or control cells were also plated together in a 24 well plate to assess growth rate of PLCG1 knockout cells relative to parental K562 Cas9 or Molm14 Cas9 cells over time. mCherry expression was detected by flow cytometry analysis on a LSRFortessa (BD Biosciences) every two days for a total of fourteen days. Cells were split every two to three days to ensure they were cultured in logarithmic growth.

RasGRP4 knockout cells: A total of 0.2×106 cells were plated in a 1mL final volume in a 24 well plate. K562 MIG EV cells (GFP positive), K562, or K562 RasGRP4 clonal cells (GFP negative) were plated alone to monitor GFP expression of logarithmic growing cells over time. An equal number (0.1×10^6^ cells) of K562 MIG EV and K562, K562 RasGRP4 knockout cells or control cells were also plated together in a 24 well plate to assess growth rate of knockout cells relative to parental K562 cells over time. GFP expression was detected by flow cytometry analysis on a LSRFortessa (BD Biosciences) every two days for a total of ten days. Cells were split every two days to ensure they were cultured in logarithmic growth.

### Cell Viability assay

Cells were exposed to imatinib or dasatinib in triplicate at 2×10^5^ cells/mL in 96 well plates (0.1mL total volume). After 48 hours, viability was assessed by CellTiter-Glo (Promega, Madison, WI, USA) on a SpectraMax M3 microplate reader. Viabilities and IC50 plots were analyzed using Prism 7 software (GraphPad, La Jolla, CA, USA).

### Caspase-3 activation assay

K562 or Molm14 cells were plated at 0.1×10^6^ cells per well of a 24-well plate and treated with DMSO (0.1%), imatinib, or dasatinib, or AC220 for forty-eight hours at 37°C. Cells were fixed with 4% paraformaldehyde for ten minutes in the dark at room temperature, washed in 1X-PBS, and permeabilized in 2mL methanol overnight at -20°C. After washing in PBS, cells were rehydrated in FACS buffer (1X-PBS supplemented with 4% fetal bovine serum) for one hour at room temperature. After centrifugation and aspiration, cells were stained with 2.5uL anti-active caspase-3 antibody conjugated to FITC (BD Biosciences) in a total volume of 50uL FACS buffer for one hour at room temperature in the dark. Cells were washed in 2mL of 1X-PBS prior to analysis by flow cytometry using a LSRFortessa machine (BD Biosciences). 20,000 ungated events were collected and the percentage of caspase-3 positive cells was analyzed on FlowJo software (Tree Star).

### Digitonin nucleotide exchange assay

Overview of assay: In this assay, cells are permeablized with digitonin to allow the entry of radioactive α-phosphate-labeled GTP. The radiolabeled GTP can be loaded onto RAS and hydrolyzed into RAS-GDP without loss of the radiolabeled nucleotide, meaning that RAS-bound radioactivity accumulates over time. Cells are subsequently lysed, RAS is immunoprecipitated, and RAS-bound nucleotide can be quantified by scintillation counting. The resulting data represents the amount of GTP loaded onto RAS as a function of time.

Experimental method: Cells were starved in phosphate-buffered saline containing 0.1% bovine serum albumin for 2 hours at 37°C, then permeabilized with 15 μg/mL digitonin and loaded with 33μCi/mL α-^32^P-GTP. Cells were lysed at the indicated time points by adding the lysis buffer (50mM HEPES pH 7.5, 100mM NaCl, 10mM MgCl_2_, 2% NP-40, 0.1% Lauryl maltoside) supplemented with protease and phosphatase inhibitors (0.1mg/mL Pefabloc, 2μg/mL leupeptin, 100 μM PMSF, 1 μg/mL pepstatin A, 100 μM sodium vanadate, 4mM β-glycerophosphate, 3.4nm microcystin), GDP and GTP (both at 100 μM) and Ras antibody Y13-259 (2.5 μg/mL). Lysates were cleared by centrifugation and Ras-antibody complexes were collected on Protein G sepharose beads. After washing, the beads were drained dry and subjected to Cerenkov counting in a Wallac 1414 Liquid Scintillation Counter for 1 minute.

## Notes

### Competing Interest Statement

The authors have declared no competing interest.

