## Supplementary Figures for "A critical role for PLCG1 in RAS activation by BCR-ABL1 and FLT3-ITD"

**Supplementary Figure 1.**

**A.** Chromatogram traces of WT K562 cells (top) and of clone K562 PLCG1-1-2 in the region of where guide RNA 1 cuts. The editing on the genomic DNA editing is highlighted.

**B.** Chromatogram traces of WT K562 cells (top) and of clone K562 PLCG1-3-10 in the region of where guide RNA 3 cuts. The editing on the genomic DNA editing is highlighted.

**Supplementary Figure 1A.**

**A. K562 sgPLCG1_1_2 TOPO Cloning Sequencing Analysis**

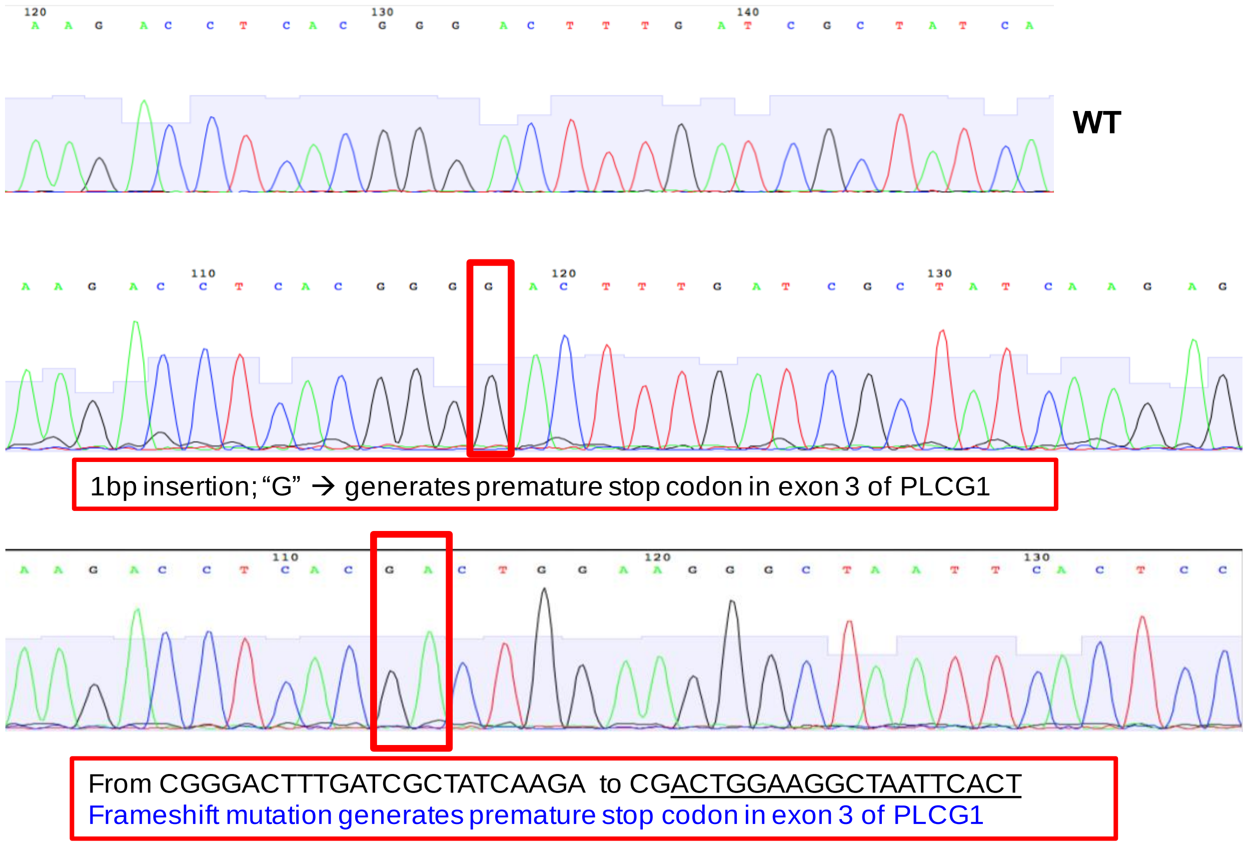

#### Supplementary Figure 1B.

**B. K562 sgPLCG1_3_10 TOPO Cloning Sequencing Analysis**

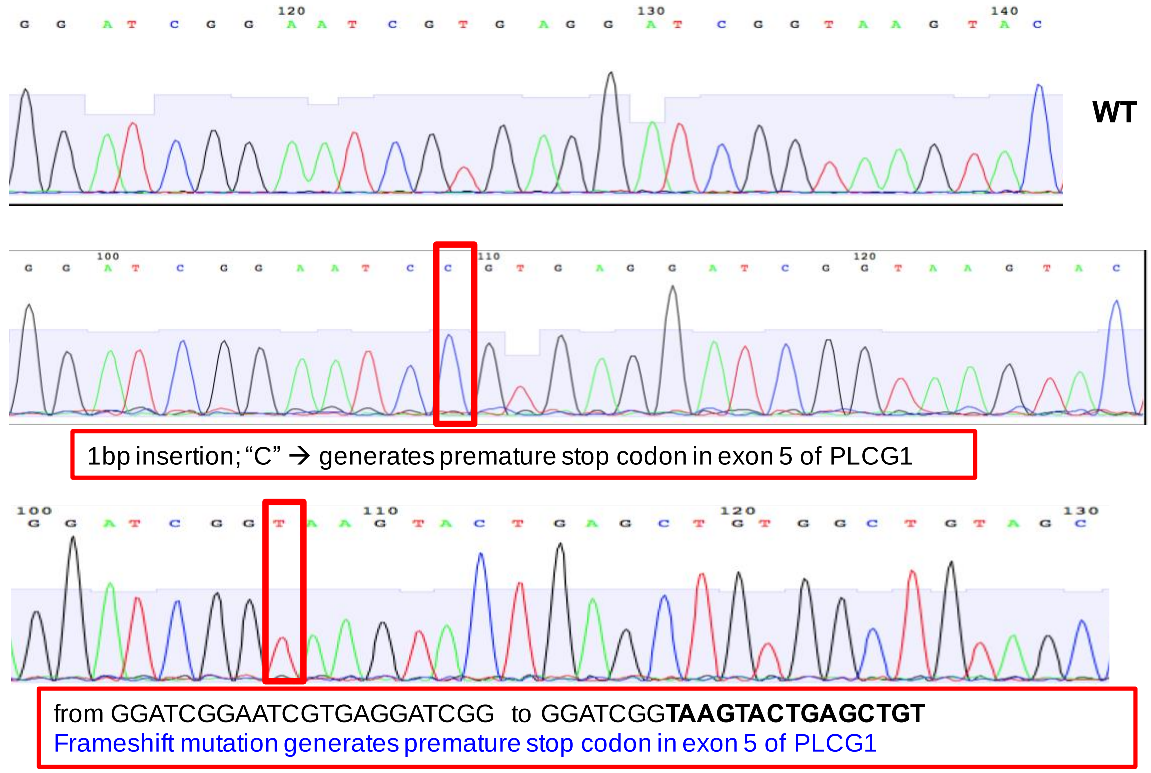

**Supplementary Figure 2. Knockout of PLCG1 decreases proliferation of BCR-ABL1 expressing cells and modulates responsiveness to BCR-ABL1 inhibitors.**

**A**. Cell viability of parental and PLCG1 knockout K562 cells was assessed with Cell Titer Glo assay following treatment with the indicated drugs of various doses for 48h.

**B**. Flow cytometric detection of cleaved caspase-3 in control and PLCG1 knockout K562 cells treated with the indicated drugs for 48h.

**C**. Addition of imatinib to PLC inhibitor decreases viability of BCR-ABL1 expressing cells.

Cell viability assessed by Cell Titer Glo assay following 48h drug treatment of K562 cells with the drugs at the indicated doses.

**D**. PLC inhibitor and BCR-ABL1 TKI is not synergistic.

Chou Talalay Combination Index (CI) was calculated from the cell viability data in (C).

**Supplementary Figure 2.**

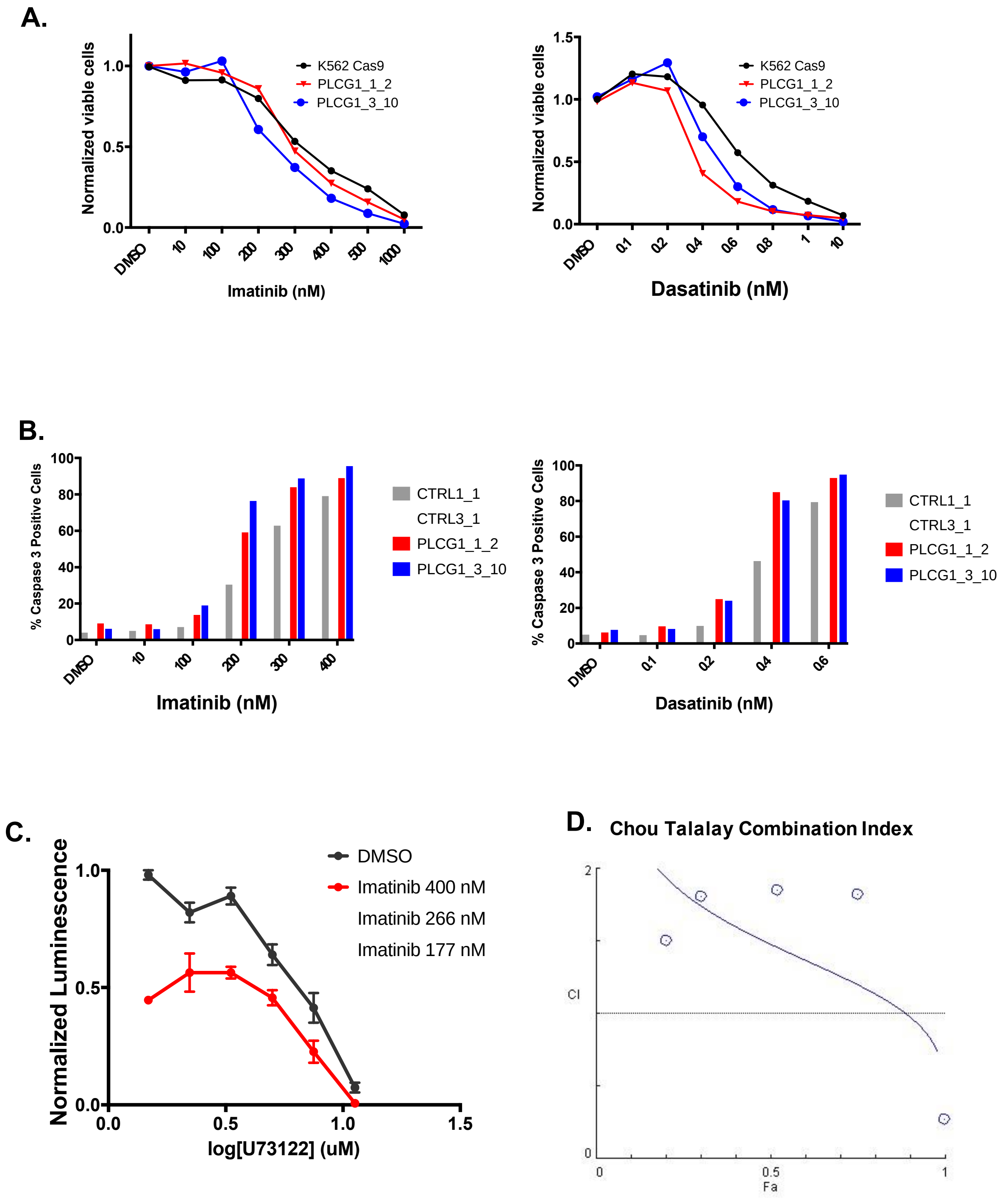

**Supplementary Figure 3. The RasGEF, RasGRP4, potentially mediates RAS activation in BCR-ABL1 expressing cells.**

**A**. DAG responsive pathway for RAS activation in BCR-ABL1 expressing cells.

Western immunoblot analysis of RAS activity in K562 cells that were serum starved for 3 hours in 0.1% RPMI followed by 25ng/mL PMA stimulation at 37˚C for the indicated time.

**B**. RasGRP4 heterozygous K562 cells have reduced RasGRP4 mRNA levels compared to parental K562 cells.

qPCR analysis of RasGRP4 mRNA levels in K562 and three different K562 RasGRP4 heterzygouslyt deleted clones via a CRISPR Cas9/gRNA ribonucleoprotein (RNP) approach.

**C**. Loss of RasGRP4 decreases growth rate of BCR-ABL1 expressing cells.

Equal number of K562 MIG EV cells (GFP positive) and K562 or RasGRP4 heterozygous deleted clonal cells (GFP negative) were plated and GFP expression was assessed over time.

**D**. Loss of RasGRP4 does not appreciably decrease Ras-GTP levels basally or following MEK inhibition.

Western immunoblot analysis of RAS, MEK, and ERK activity in cells treated with DMSO, MEK inhibitor (500nM PD0325901), or MEK inhibitor and Imatinib (1uM) for 12 hours.

### Supplementary Figure 3.

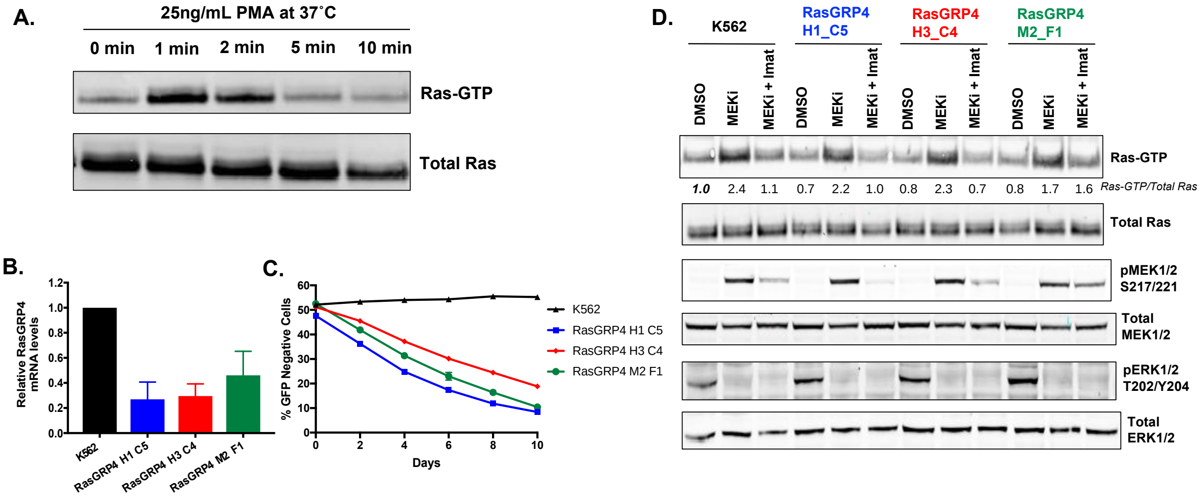
